# Regulation of Macrophage IFNγ-Stimulated Gene Expression by the Transcriptional Coregulator CITED1

**DOI:** 10.1101/2022.08.16.504142

**Authors:** Aarthi Subramani, Maria E. L. Hite, Hursha Kondee, Grace E. Millican, Erin E. McClelland, Rebecca L. Seipelt-Thiemann, David E. Nelson

## Abstract

Macrophages are highly dynamic innate immune cells that adopt temporary phenotypes, known as polarization states, in response to changing environmental signals to combat microbial infection and contribute to the maintenance of tissue homeostasis. During the early stages of an infection, exposure to interferon-gamma (IFNγ) and microbial ligands induce M1 polarization, a heightened proinflammatory anti-microbial state, by regulating the expression of interferon stimulated genes (ISGs). While this response must be sufficiently vigorous to ensure the successful clearance of invading pathogens, it must also be spatially and temporally restricted to prevent uncontrolled inflammation that could result in tissue damage and disease. This is controlled by a variety of cell-extrinsic and -intrinsic mechanisms, including the expression of CBP/p300-interacting transactivator with glutamic acid/aspartic acid-rich carboxyl-terminal domain 2 (CITED2), a transcriptional coregulator that limits IFNγ-stimulated proinflammatory gene expression by inhibiting STAT1- and IRF1-regulated ISGs. In this study, we show that CITED1, another member of the CITED family of proteins, is itself an ISG and is expressed in a STAT1-dependent manner, and that IFNγ stimulates the nuclear accumulation of fluorescently tagged CITED1 proteins. In contrast to CITED2, ectopic expression of CITED1 enhanced the expression of a subset of ISGs, including *Ccl2, Ifit3b, Isg15*, and *Oas2*. This effect was reversed in a *Cited1* null cell line produced by CRISPR-based genomic editing. Collectively, these data show that CITED1 helps to maintain proinflammatory gene expression during periods of prolonged IFNγ exposure and suggests a distinct and antagonistic relationship between CITED proteins in the regulation of macrophage inflammatory function.

## Background

Macrophages are multifunctional innate immune cells that play a major role in the maintenance of tissue homeostasis (1). These tissue-associated phagocytic cells respond rapidly to injury and infection, initiating an appropriate inflammatory response and clearing microbial pathogens and cellular debris before participating in wound-healing and attenuating inflammatory signals as the infection is resolved. As part of this process, macrophages detect and respond to microbial ligands using pattern recognition receptors (PRRs), including the toll-like receptor family (TLRs) of proteins. These stimulate the activity of nuclear factor kappa-light-chain-enhancer of activated B cells (NF-κB) and other proinflammatory transcriptional regulators, that increase the expression of cytokines, chemokines, and anti-microbial proteins, such as inducible nitric oxide synthase (iNos) (2-4). This response is modified or enhanced by a variety of endogenous factors, including interferon-gamma (IFNγ), which is largely produced by T cells and natural killer cells (5).

IFNγ, also known as type II interferon, is a pleotropic cytokine capable of regulating the activity of innate and adaptive immunity, and is associated with the response to viral and non-viral pathogens (6). Within the context of innate immunity, IFNγ stimulates classical activation or M1 polarization of macrophages (7), a heightened proinflammatory anti-microbial state that is important for the successful clearance of bacterial and eukaryotic pathogens, including the fungal pathogen, *Cryptococcus neoformans* (8-11). This transition to the M1 state is accompanied by extensive transcriptional reprogramming, involving over 1000 genes (12,13). These changes in gene expression are largely directed by the transcription factors signal transducer and activator of transcription 1 (STAT1) and interferon regulated factor 1 (IRF1), operating individually or in concert at IFNγ-stimulated gene (ISG) promoters (14).

IFNγ homodimers stimulate STAT1 activity by binding to and facilitating the assembly of tetrameric interferon-gamma receptor (IFNGR) complexes from dimers of IFNGR1 and IFNGR2. This activates receptor-associated Janus kinase (JAK) 1 and 2 by transphosphorylation within the cell, which subsequently tyrosine-phosphorylate the cytosolic domain of IFNGR1. This enables STAT1 proteins to dock with the receptor complex via phospho-tyrosine-binding Src-homology-2 (SH2) domains, bringing these transcription factors into proximity with the activated JAK proteins, which phosphorylate Y701 within the STAT1 C-terminus. The same SH2-domains used for receptor-binding also facilitate the homodimerization of phosphorylated STAT1 proteins, creating γ-activated factors (GAF). These GAFs translocate to the nucleus and initiate a first phase of ISG expression through binding to STAT1 target promoters that contain palindromic γ-activated sites (GAS) (15). This includes the *Irf1* gene and newly synthesized IRF1 proteins then participate in the regulation of a second wave of gene expression through binding interferon-stimulated response elements (ISRE) and IRF-response elements in ISG promoters (16,17). These include interferon-induced protein with tetratricopeptide repeat 1 (*Ifit1)*, interferon-stimulated gene 15 (*Isg15*), MX dynamin like GTPase 1 (*Mx1*), and 2’-5’-Oligoadenylate synthase 1 (*Oas1*) (14). Additionally, IRF1 and STAT1 co-regulate genes that contain both GAS and ISRE cis-regulatory sites (18-20). This includes *Irf9*, bone marrow stromal cell antigen 2 (*Bst2*), and interferon induced transmembrane protein 1 (*Ifitm1*) (21-23), but also *Stat1* itself, constituting a positive feedback loop (24).

To prevent tissue damage that accompanies uncontrolled or prolonged inflammation, macrophage IFNγ signaling is restrained by a variety of mechanisms (25-27). It is antagonized by anti-inflammatory cytokines, interleukin-4 (IL-4) and IL-13, which promote an alternative activation or M2 polarization state and the downregulation of many ISGs. Negative feedback also occurs through expression of cell-intrinsic factors, including suppressor of cytokine signaling 1 (Socs), an ISG co-regulated by STAT1 and IRF1 that functions as a potent inhibitor of IFNγ signaling at the JAK level (28-31). More recently, CBP/p300-interacting transactivator with glutamic acid (E) and aspartic acid (D)-rich tail 2 (CITED2) has been identified as a transcriptional co-regulator that operates at the chromatin/promoter level to attenuate macrophage proinflammatory gene expression (32).

CITED2 is one of three CITED family proteins present in mammalian systems (CITED1, 2, and 4(33-36)). Unable to bind directly to DNA, CITED proteins increase or inhibit the expression of genes by facilitating or preventing transcription factors from forming chromatin complexes with the histone acetyltransferase, CREB-binding protein (CBP) or its paralog p300. These interactions require a C-terminal conserved region 2 (CR2) domain common to all CITED family proteins (37), and an unstructured N-terminal region that differs between CITED proteins. While the CR2 domains binds cysteine-histidine (CH; also known as TAZ) domains within CBP/p300, the unique N-terminus of each CITED family protein facilitates interactions with different sets of transcription factors. In this way, CITED proteins operate as adaptors, stabilizing transcription factors:CBP/p300 complexes. However, they can also blocking the formation of other complexes competitively in instances where transcription factors interact with CBP/p300 via the same CH domain as the CITED protein. For example, CITED2 is constitutively expressed in myeloid cells, including macrophages, and localizes exclusively to the nucleus in complex with CBP/p300 (38). Here, it restricts the ability of hypoxia-inducible factor 1 alpha (HIF1α) and p65-containing NF-κB transcription factors to access the CH1 domain of CBP/p300, thereby reducing proinflammatory gene expression (38-42). In fact, synthetic constrained peptides derived from the CR2 domain of CITED2 function as potent inhibitors of HIF1α signaling (43). CITED2 has also been shown to repress both STAT1- and IRF1-dependent ISGs in bone-marrow-derived macrophages and RAW264.7 macrophage-like cells, likely by a similar mechanism (44).

Until recently, CITED2 was thought to be the only CITED family member expressed in macrophages (32). In this study, we show that IFNγ stimulation promotes expression of *Cited1* in a STAT1-dependent manner and nuclear accumulation of CITED1 proteins. Unlike CITED2, expression of ectopic CITED1 largely enhanced the expression of STAT1- and IRF1-dependent ISGs in IFNγ-stimulated macrophages, and this was reversed in *Cited1* null cells. As CITED1 expression is featured as part of a later wave of ISGs, these data indicate that it may serve as a mechanism to prolong the IFNγ response for a subset of STAT1- and IRF1-regulated genes.

## Results

### Cited1 and 2 respond differently to M1-polarizing stimuli

Over the past few years, *Cited2* has been found to play an important role in innate immune function (32,41,42,44,45). Highly expressed at both the transcript and protein level in monocytes and macrophages of murine and human origin (32), it is vital for the development of these cells (46,47), and attenuates proinflammatory gene expression by acting as an inhibitor of NF-κB, HIF1α, STAT1, and IRF1 transcriptional regulators (32,42,44). In contrast, expression of the two other mammalian *Cited* family members, *Cited1* and *4*, were documented as either undetectable or ∼100-fold lower than *Cited2*, which suggest they have no biological role in macrophages (32). However, in our previous study, we showed that infection of M1 polarized RAW264.7 murine macrophages with *C. neoformans* promotes transcriptional upregulation of *Cited1* (48). A more detailed reanalysis of these data comparing vehicle and IFNγ-treated cells showed that IFNγ stimulation alone was sufficient to induce *Cited1* (Fig. 1A), raising the possibility that *Cited1* expression was a feature of M1 polarization and *Cited1* might also be an ISG.

**Figure 1:**
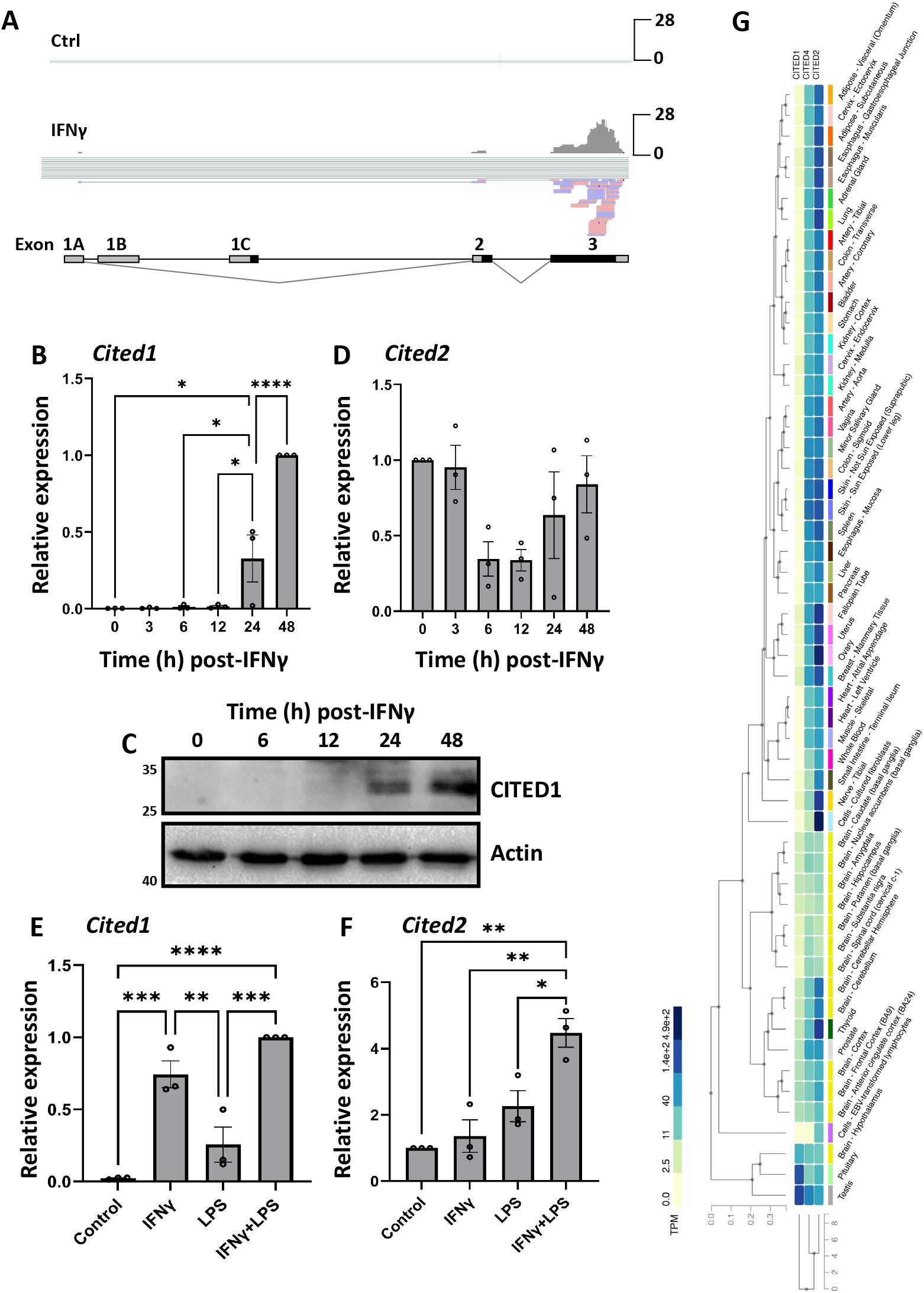
*Cited1* is an IFNγ-responsive gene. (A) RNAseq analysis was performed on mRNA extracted from RAW264.7 cells treated +/- IFNγ for 24 h. Reads were aligned to the *Cited1* gene using Integrated Genome Viewer. Protein coding regions of exons are marked in black. An individual representative repeat (1 of 3) is shown for each condition. (B-D) RAW264.7 cells were incubated with IFNγ for the indicated times or (E+F) IFNγ and/or LPS for 24 h then RNA and proteins were harvested. (B+D-F) Expression of *Cited1+2* transcripts was measured by qRT-PCR. Error is represented as S.E. Statistical differences between samples were appraised using one-way analysis of variance (ANOVA) followed by a Tukey’s multiple comparison test. Statistical significance is indicated as follows: *, p<0.05; **, p<0.01; ***, p<0.001; ****, p<0.0001. (C) CITED1 protein levels were measured by western blotting. (G) GTEx analysis of *Cited* family member expression across 54 different non-diseased tissues.

To validate this result and investigate the dynamics of IFNγ-stimulated *Cited1* expression, RNA was harvested from RAW264.7 cells at 3, 6, 12, 24, and 48 h post-IFNγ treatment and *Cited1* transcript levels were measured using qRT-PCR. Here, *Cited1* levels were apparent at 24 h, and were further increased by 48 h post-stimulation (>300-fold increase in t=0 vs. 48 h; Fig. 1B), and this was mirrored at the protein level (Fig. 1C). This differed from the reported effects of M1 polarizing stimuli on *Cited2* expression, which has been shown to be repressed by IFNγ or lipopolysaccharide (LPS) at 6 h post-treatment (32). We also observed a similar decrease in *Cited2* transcript levels was observed post-IFNγ, but this was transient and not statistically significant, with *Cited2* returning to basal levels within 24 h. To explore the effects of other M1-polarizing stimuli on *Cited1* and *2* expression, macrophages were treated with LPS alone and in combination with IFNγ. Here, LPS had no effect on *Cited1* expression and did not enhance IFNγ-stimulated expression of the gene (Fig. 1E). In contrast, LPS or IFNγ treatment alone had no effect on *Cited2* expression at 24 h post-stimulus (Fig. 1F). However, IFNγ and LPS co-treatment stimulated a >4-fold increase in *Cited2* expression compared to vehicle. These contrasting expression patterns indicate that *Cited1* and *2* are regulated differently and likely play distinct roles in modulating the macrophage transcriptome, as each operates on a differing timescale and in a stimulus-dependent fashion.

To further explore the notion that regulation of *Cited1* and 2 differ, the expression of *Cited* family genes was examined using the open access Genotype-Tissue Expression (GTEx) database, which contains searchable gene expression data based on the molecular analysis of 54 non-diseased human tissue sites from ∼1000 individuals (49). This showed that the expression of *Cited1* and *Cited2* was strikingly different. Whereas *Cited2* expression was near-ubiquitous, expressed in most tissues, *Cited1* was largely restricted to the testis and pituitary (Fig. 1G). These differences are corroborated by experimental data showing that although both *Cited1* and *2* are expressed in the developing kidney, only *Cited2* persists in mature renal structures (50). Collectively, these data indicate that *Cited1* and *2* are regulated differently and are unlikely to be functionally redundant in macrophages and other contexts.

### Regulation of Cited1 expression by STAT1

While differences in transcriptional regulation of the murine *Cited1* and *Cited2* gene is now clear, the finer detail of *Cited1* gene expression is lacking. Prior studies have shown that the 1.0 kb region immediately upstream of the TATA box has promoter activity and contains putative binding sites for Sp1, Oct-1, and AP-2, although none of these have been experimentally confirmed (51). From these data, it is unclear how IFNγ stimulates *Cited1* expression in macrophages.

While engagement of IFNγ-receptors activates a range of signaling pathways (52), the JAK:STAT pathway is considered the central coordinator of the transcriptional response with STAT1 transactivating ISGs directly or via IRF1 (53,54), which is itself a STAT1-regulated gene (15,55) (Fig. 2A). To investigate whether *Cited1* is regulated by the STAT1-IRF1 axis, we scanned a region spanning -2000 to 100 bp relative to the transcriptional start site of the murine *Cited1* gene using the Eukaryotic Promoter Database (Swiss Institute of Bioinformatics) for transcription factor binding sites (56,57). A total of 3 STAT1 (-18, -724, and -1038) and 2 IRF1 (-1237, and - 1297) putative cis-regulatory sites were identified (cut-off p-value <0.001; Fig. 2B). Based on these data, we hypothesized that IFNγ-stimulated *Cited1* expression was regulated by the STAT1-IRF1 axis and was therefore STAT1-dependent.

**Figure 2:**
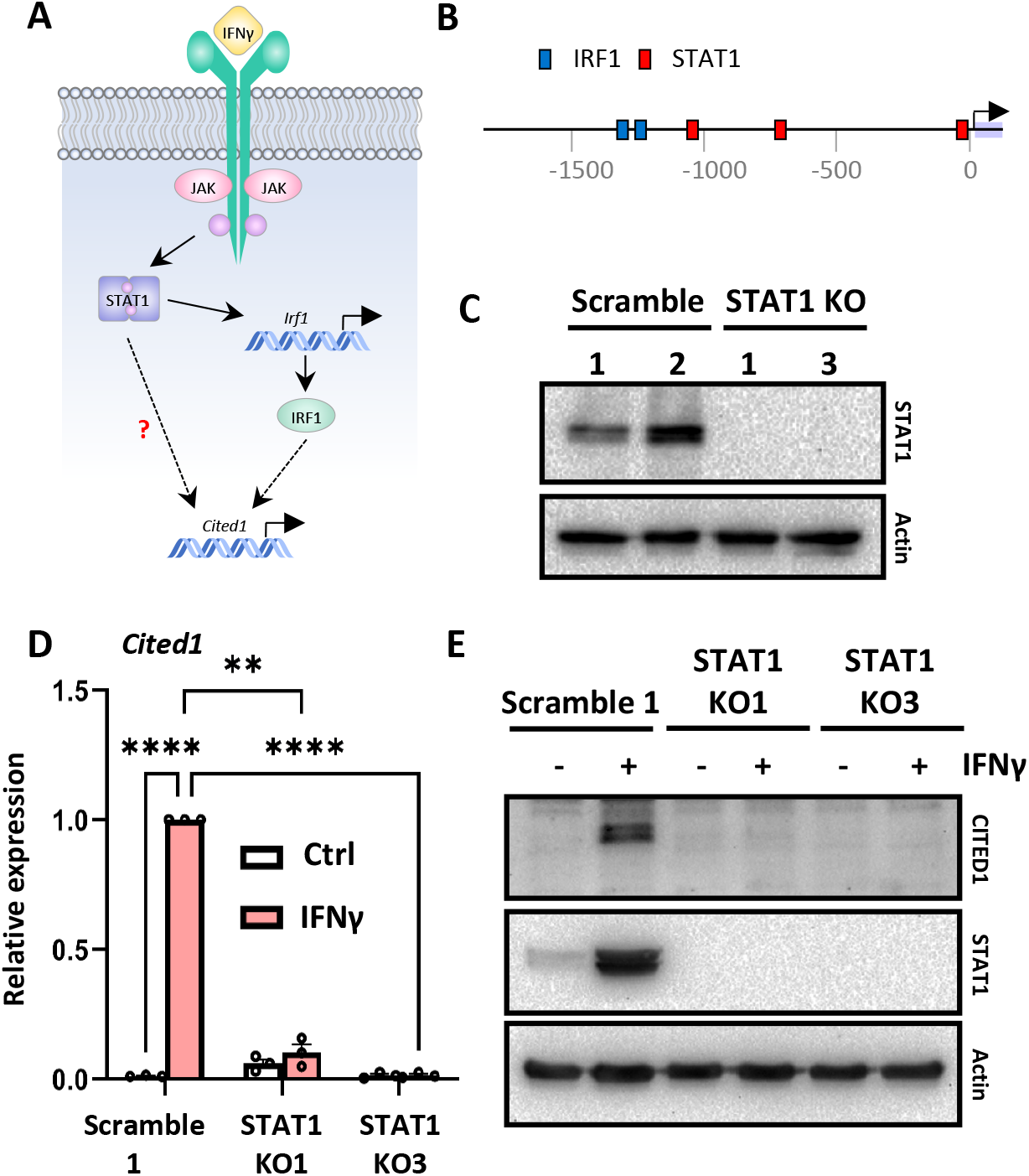
*Cited1* expression is STAT1-dependent. (A) Diagram of IFNγ-STAT1 signaling pathway. The relationship under investigation is demarcated by a question mark. (B) Putative STAT1 and IRF1 transcription factor binding sites were detected in the murine *Cited1* promoter using EPD and a p-value cutoff of ≤0.001. (C) Western blot analysis for STAT1 proteins in two independent clonal RAW264.7 cell lines stably transduced with lentiviral constructs to express Cas9 and either non-targeting (scramble) or *Stat1*-targeting gRNA. (D+E) RAW264.7 STAT1 KO cells and scramble controls were stimulated with IFNγ for 48 h. (D) *Cited1* expression was measured by qRT-PCR and (E) CITED1 proteins were detected by western blotting. Error is represented as S.E. Statistical differences between samples were appraised using one-way ANOVA followed by a Tukey’s multiple comparison test. Statistical significance is indicated as follows: **, p<0.01; ****, p<0.0001.

To test this, RAW264.7 cells were transduced with a lentiviral construct to express the Cas9 endonuclease and either a non-targeting ‘scramble’ guide RNA (gRNA) or a gRNA targeting exon 9 of the *Stat1* gene, clonal lines were produced, and loss of STAT1 protein expression was examined by western blot (Fig. 2C). As predicted, IFNγ-stimulated *Cited1* RNA expression was abrogated in both STAT1 null clonal lines but remained robust in the scramble controls (∼80-fold increase; Fig. 2D). This result was confirmed at the protein level where CITED1 was detected in IFNγ-stimulated scramble control cells but not STAT1 null cells (Fig. 2E). The increased STAT1 expression observed in the control cells is consistent with the known effects of IFNγ priming on STAT1 expression (58,59). Collectively, these data indicated that the *Cited1* gene was downstream of the JAK:STAT portion of the pathway and was regulated by STAT1 transcription factors, although it is unclear if STAT1 directly regulates *Cited1* or *Cited1* resides further down the pathway.

### IFNγ stimulated nuclear translocation of CITED1

To function as a transcriptional co-regulator, CITED1 must be present in the nucleus. However, its subcellular localization varies among cell types and is cytosolic in most, likely due to a nuclear export sequence present within the CR2 domain (60,61). To determine the localization of the protein in macrophages, RAW264.7 cells were stably transduced with pINDUCER20-EYFP-CITED1 to express full-length murine CITED1 with an N-terminal enhanced yellow fluorescent protein (EYFP) tag under the control of a doxycycline (dox)-dependent promoter. Cells were incubated with dox to induce EYFP-CITED1 prior to cytokine stimulation. Prior to IFNγ treatment, EYFP-CITED1 was predominantly cytosolic in all cells (Fig. 3A+B). However, IFNγ stimulated the relocalization of the protein, with EYFP-CITED1 becoming enriched in the nucleus by 24 h post-stimulation and remaining there for at least a further 24 h post-treatment (Fig. 3A+B).

**Figure 3:**
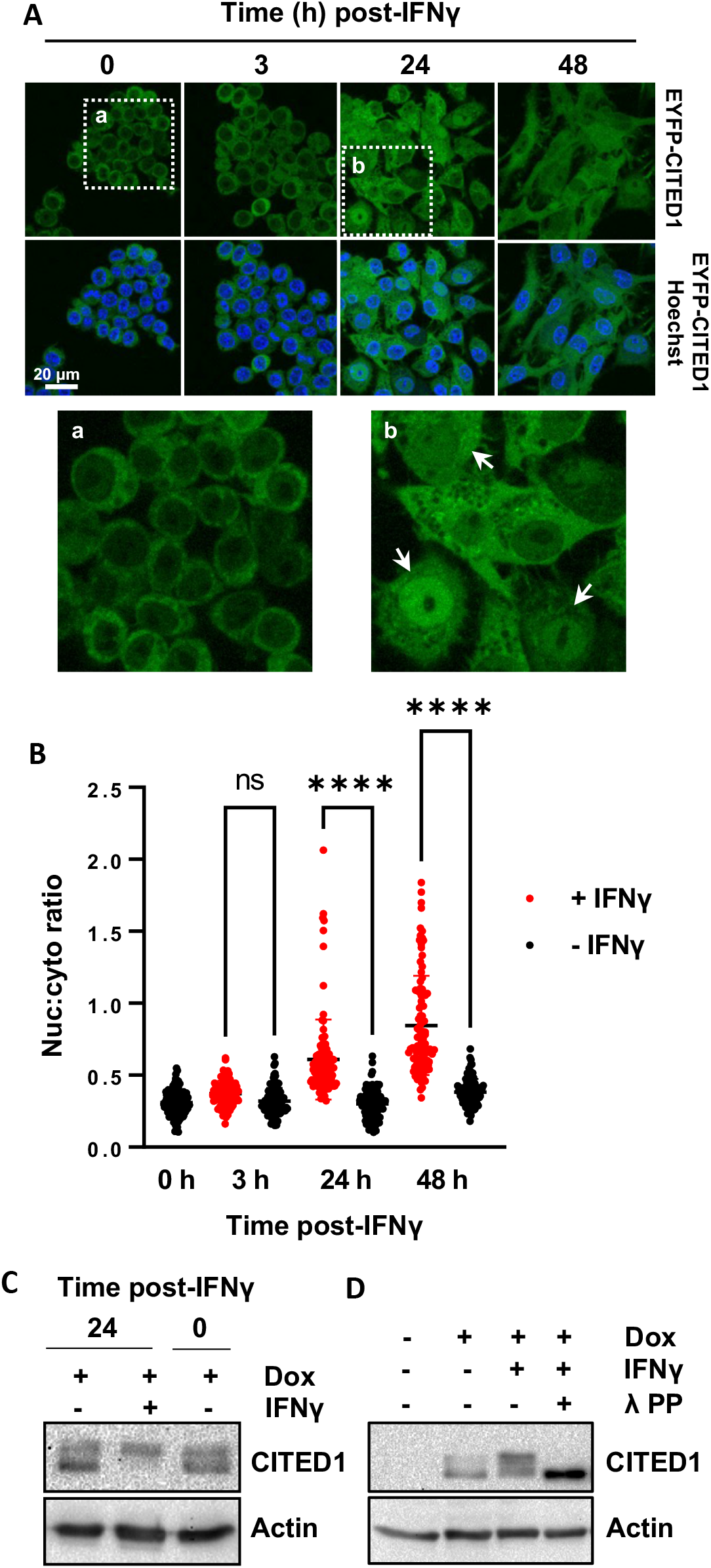
IFNγ stimulates nuclear translocation and phosphorylation of CITED1. (A) RAW264.7 stably transduced with lentiviral constructs to express EYFP-CITED1 from a dox-inducible promoter (DI-EYFP-CITED1 cells). Cells were incubated with dox for 24 h prior to co-treatment with IFNγ for the indicated times and visualized by live cell confocal microscopy. Arrow heads mark cells where nuc:cyto EYFP-CITED1 ≥ 1. (B) Quantification of nuc:cyto EYFP-CITED1 in individual cells from the experiment described in (A). Data are plotted for ≥100 cells/condition. Statistical differences between samples were appraised using one-way ANOVA followed by a Tukey’s multiple comparison test. Statistical significance is indicated as follows: ****, p<0.0001. (C) DI-CITED1 cells were incubated with dox for 24 h to stimulate CITED1 expression prior to co-treatment with IFNγ or vehicle for a further 24 h. Low- and high-mobility CITED1 species were detected by western blotting. (D) DI-CITED1 cells were treated as described in (C) but lysed at 6 h post-IFNγ in a non-denaturing 1% triton X-100 buffer. Lysates were incubated +/- λ protein phosphatase for 30 min prior to western blot analysis for CITED1.

Phosphorylation has also been shown to affect CITED1 localization. In MC3T3-E1 murine osteoblasts, parathyroid hormone (PTH) has been shown to stimulate the expression and nuclear translocation of CITED1 proteins as part of the osteoblastic differentiation process (62,63). Here, the nuclear accumulation of CITED1 required the PKC-dependent phosphorylation of the protein at Ser79 (62). To test whether phosphorylation of CITED1 accompanied IFNγ-induced nuclear translocation, we generated RAW264.7 that expressed untagged CITED1 protein in a dox-dependent manner (DI-CITED1). This decoupled CITED1 expression from IFNγ stimulation and allowed cells to produce a preexisting pool of exogenous CITED1. The phosphorylation state of these proteins were measured pre- and post-IFNγ treatment. While there are currently no antibodies available to directly detect phosphoryated forms of the protein, phosphorylation has been shown to reduce the mobility of CITED1 on SDS-PAGE gels (60). Prior to IFNγ treatment, CITED1 appeared as two major bands, a high mobility band and a fainter low mobility band (Fig. 3C). However, in cells incubated with IFNγ for 24 h, only the low-mobility CITED1 band was apparent, indicating that the cellular pool of CITED1 proteins transition from a largely dephosphorylated state to a mostly phosphorylated state upon treatment with IFNγ. The appearance of the higher molecular weight band was eliminated by λ protein phosphatase treatment (λPP) of the samples prior to western blotting, confirming that this band was associated with phosphorylated forms of CITED1 and was not caused by other post-translational modifications (Fig. 3D). Collectively, these data raise the possibility that IFNγ regulates the localization of CITED1 in a phosphorylation-dependent manner, similar to parathyroid hormone-stimulated nuclear translocation of CITED1 in osteoblasts.

### CITED1 enhanced the expression of a subset of ISGs

As IFNγ treatment promoted nuclear accumulation of CITED1, we speculated that it may also function as a transcriptional co-regulator under these conditions. CITED2, which is constitutively nuclear, inhibits proinflammatory gene expression by competing with NF-κB and HIF-1 transcription factors for binding of the CH1 domain of CBP/p300 (38,41). Additionally, and of particular relevance to this study, overexpression of CITED2 in RAW264.7 macrophages inhibits IFNγ-stimulated expression of genes regulated by the STAT1-IRF1 axis, including *Irf1, Irg1, F3, Tmem140*, and *Dnase13* (44). While, CITED1 is also known to interact with CBP/p300, the two CITED family proteins exhibit differing binding preferences for the CH domains in CBP/p300; CITED1 only weakly interacts with the CH1 domain and instead shows a preference for the central CH2 domain (37), distant from known STAT1 binding regions (64). For this reason, we hypothesized that CITED1 may have distinct effects on STAT1- and IRF1-dependent gene expression. To explore this, the DI-CITED1 cell line was utilized as part of a gain-of-function strategy, measuring the effect of ectopic CITED1 expression on IFNγ-stimulated gene expression. In brief, gene expression changes were measured 24 h post-IFNγ by RNAseq-based transcriptome profiling with and without prior dox incubation (Fig. 4A). Corresponding control samples were prepared using unmodified RAW264.7 to identify and exclude gene expression changes stimulated by dox alone.

**Figure 4:**
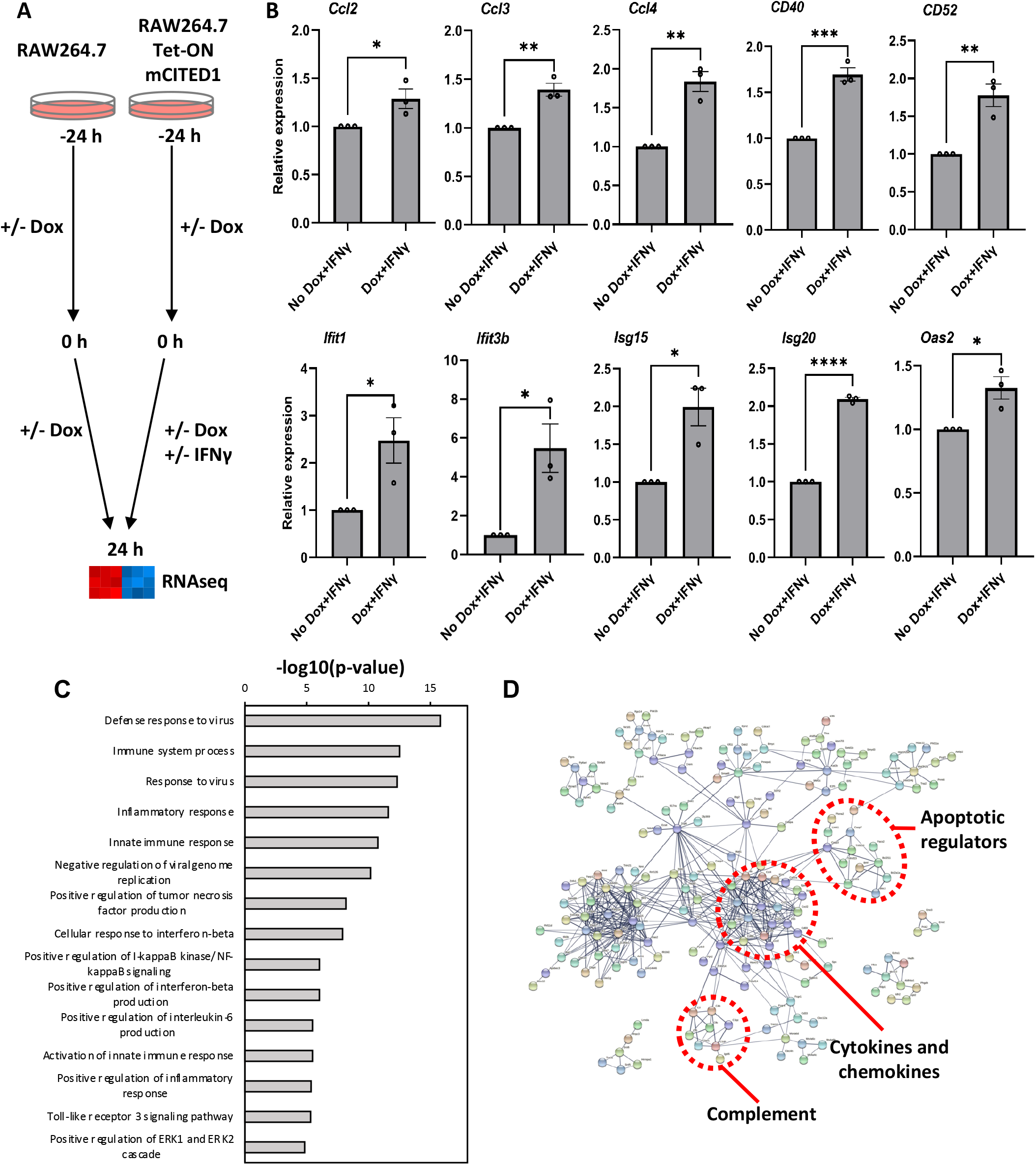
CITED1 enhances IFNγ-stimulated gene expression. (A) Design of CITED1 overexpression transcriptome profiling experiment. RAW264.7 DI-CITED1 cells were incubated +/- dox for 24 h to over-express CITED1 prior to co-treatment with IFNγ or vehicle for a further 24 h. Following treatments, total RNA was harvested for RNAseq. (B) Select CITED1-regulated genes identified by the RNAseq screen were validated by qRT-PCR. (C) GO analysis was performed in DAVID on DEGs from a pairwise comparison of DI-CITED1 cells treated with -dox +IFNγ or +dox +IFNγ (IFNγ:dox+IFNγ). Top biological process (BP) GO terms are ranked by - log(*p*-value). (D) STRING analysis was performed on the same set of DEGs. Boundaries enclosing gene clusters denote gene sets with a common function, as identified using the KEGG analysis tools in STRING.

An initial assessment of the transcriptome data performed using a CuffDiff-based analysis pipeline identified 724 differentially expressed genes (DEGs) in a pairwise comparison of IFNγ-stimulated cells with and without dox treatment (IFNγ vs. dox+IFNγ; Table 1). These data showed increased expression of multiple members of gene families closely associated with the response to IFNγ in CITED1 over-expressing cells, including members of the C-C motif chemokine ligand (*Ccl; Ccl2, Ccl3, Ccl4, and Ccl7*), *Ifit* (*Ifit1, Ifit3, and Ifit3b*), and *Isg* (*Isg15* and *Isg20*) gene families. A selection of these were validated by qRT-PCR together with *Cd40* and *Cd50*, which encode surface markers that were also upregulated by CITED1 over-expression (Fig. 4B). Notably, increased expression of multiple genes that were up-regulated in CITED2 knockout bone marrow-derived macrophages (BMDMs) were observed, including *Mxd1, Il17ra, Tmem140, CD86*, and *Scimp (44)*, supporting the notion that CITED1 and 2 had opposing effects on IFNγ-stimulated gene expression.

**Table 1:** Selected ISGs affected by CITED1 expression.

| Cited1 O/E |  |  |  | Cited1 KO |  |  |
| --- | --- | --- | --- | --- | --- | --- |
| Gene | FC | Direction | *q*-value | FC | Direction | *q*-value |
| Cited1 | 7.55 | UP | 0.000387 | 6.71 | DOWN | 0.000364 |
| Ccl2 | 2.32 | UP | 0.000387 | 39.3 | DOWN | 0.000364 |
| Ccl3 | 2.4 | UP | 0.000387 | 40.96 | DOWN | 0.000364 |
| Ccl4 | 2.5 | UP | 0.000387 | 22.97 | DOWN | 0.000364 |
| Ccl7 | 2.18 | UP | 0.011804 | 70.93 | DOWN | 0.002853 |
| Cd40 | 2.78 | UP | 0.000387 | 2.65 | DOWN | 0.000364 |
| Cd52 | 2.12 | UP | 0.000387 | 2.91 | DOWN | 0.000364 |
| Ifit1 | 5.11 | UP | 0.000387 | 15.57 | DOWN | 0.000364 |
| Ifit3 | 5.43 | UP | 0.000387 | 18.77 | DOWN | 0.000364 |
| Ifit3b | 15.97 | UP | 0.000387 | 49.77 | DOWN | 0.000364 |
| Isg15 | 3.99 | UP | 0.000387 | 7.51 | DOWN | 0.000364 |
| Isg20 | 2.75 | UP | 0.000387 | 3.02 | DOWN | 0.000364 |
| Mx1 | 4.95 | UP | 0.000387 | 8.77 | DOWN | 0.000364 |
| Mx2 | 2.49 | UP | 0.000387 | 4.73 | DOWN | 0.000364 |
| Oas2 | 3.28 | UP | 0.000387 | 9.14 | DOWN | 0.000364 |
| Tlr9 | 2.45 | UP | 0.000387 | 2.24 | DOWN | 0.000364 |
| Tmem140 | 2.16 | UP | 0.000387 | 10.72 | DOWN | 0.000364 |

Top biological process (BP) gene ontology (GO) terms associated with DEGs from the IFNγ vs. dox+IFNγ pairwise comparison included ‘Defense response to virus’, ‘Innate immune response’, and ‘Inflammatory response’, indicative that CITED1 expression affected the expression of gene sets fundamental to the IFNγ response (Fig. 4C). Protein interaction network analysis was performed in STRING, which also allowed for the visualization of changes in the IFNγ response stimulated by CITED1 expression (Fig. 4D). Here, genes clusters associated with cytokines and chemokines (GO term ‘Regulation of cytokine production’; false discovery rate (FDR) 2.57×10^−9^), apoptosis (GO term ‘Regulation of apoptotic process’; FDR 3.9×10^−4^) and a smaller cluster of genes associated with the complement system were observed.

To more formally assess the impact of CITED1 expression on the transcriptional changes that accompany IFNγ-stimulated M1 polarization, the IFNγ vs. dox+IFNγ DEG set was compared to corresponding DEGs from a comparison of untreated control and IFNγ-treated cells (Ctrl vs. IFNγ). Of the 2775 DEGs between control and IFNγ-stimulated cells, 300 (10.8%) were affected by CITED1 overexpression (Fig. 5A), which corresponded to 41.5% of all DEGs from the IFNγ vs. dox+IFNγ pairwise comparison. Of those genes common to both pairwise comparisons, the majority (255; 85%) were concordant (Fig. 5B), indicating that CITED1 expression enhanced the effect of IFNγ stimulation on a subset of IFNγ-responsive genes. These common concordant genes included *Isg15, Isg20*, various *Ifit* family genes, and the cytokines *Ccl2* and *Ccl4* (Table 2). GO analysis of this common pool of DEGs revealed an enrichment in genes associated with macrophage antiviral function (Fig. 5C), as indicated by the GO terms ‘Defense response to virus’ (p=6.84×10^−17^; FDR=1.24×10^−13^) and ‘Negative regulation of viral genome replication (p=7.49×10^− 10^; FDR=2.72×10^−7^). Overall, these data show that CITED1 expression potentiates the effects of IFNγ stimulation on select ISGs.

**Table 2:**
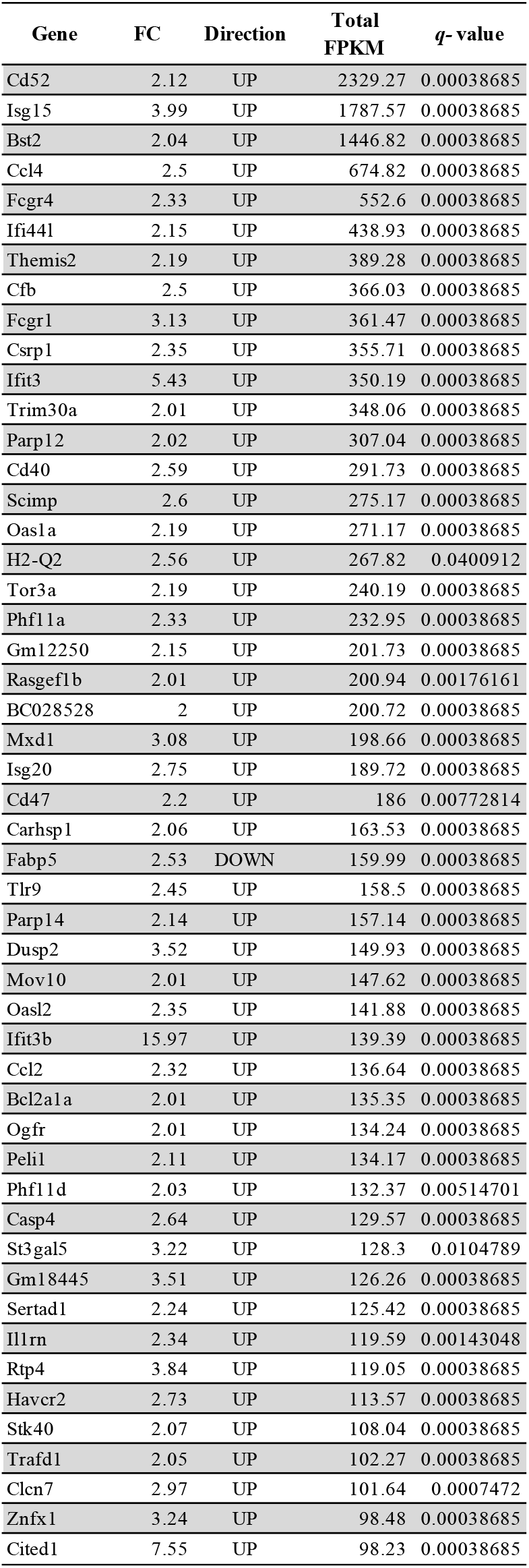
Top 50 genes (by FPKM) from IFNγ:dox+IFNγ common to Ctrl:IFNγ.

**Figure 5:**
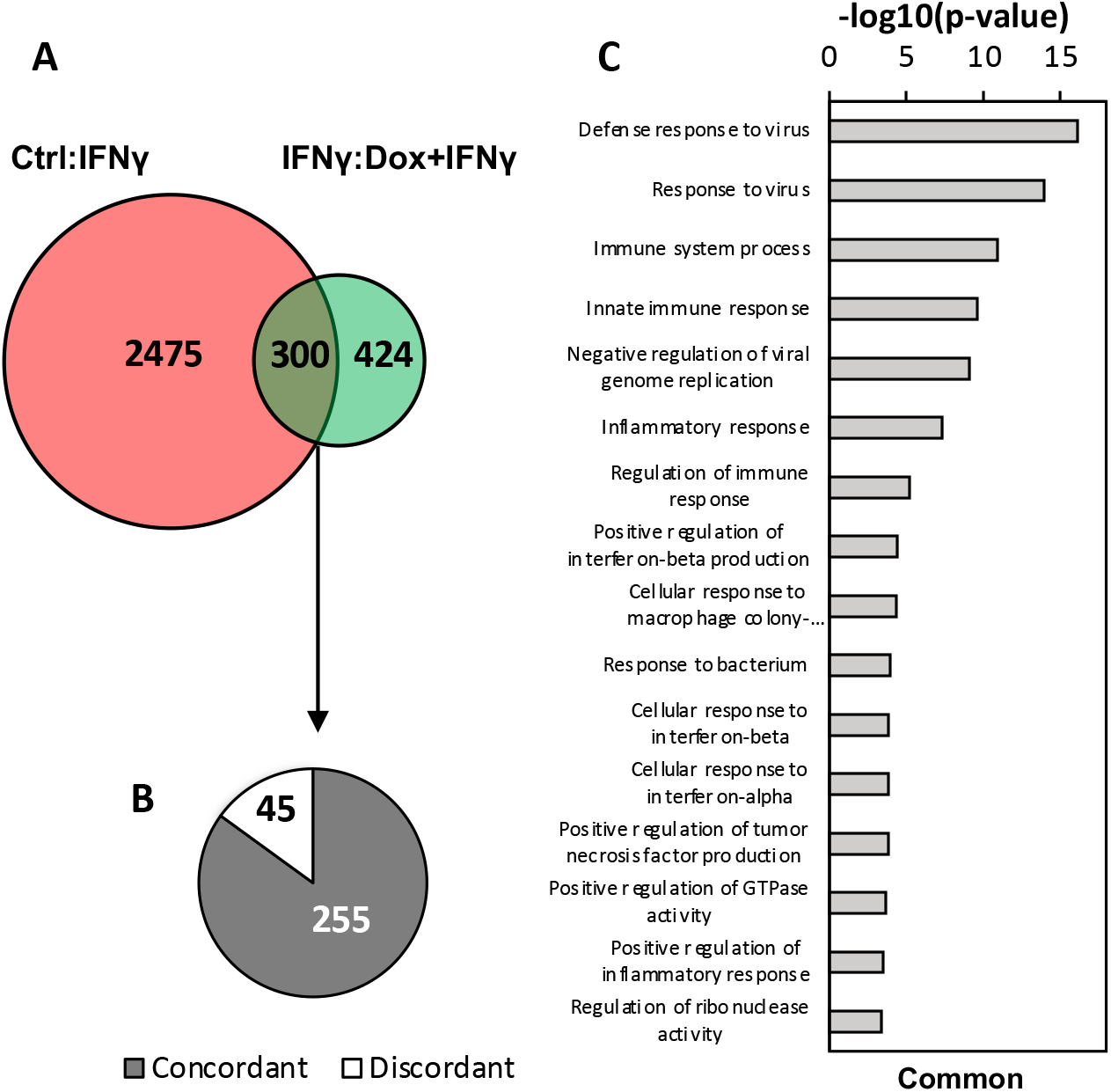
Effect of CITED1 expression on the M1 transcriptome. (A) Venn diagram to represent common and DEGs between pairwise comparisons of DI-CITED1 cells treated as follows; -dox -IFNγ:-dox +IFNγ (Ctrl:IFNγ) and IFNγ:dox+IFNγ. (B) Of the 300 DEGs common to Ctrl:IFNγ and IFNγ:dox+IFNγ 255 (85%) were concordant (grey) and 45 (15%) were discordant. (C) GO analysis was performed in DAVID on common DEGs from the comparison shown in (A). Top biological process (BP) GO terms are ranked by -log(*p*-value).

### CITED1 modulated the expression of STAT1 and IRF1 target genes

As an alternative method to examine the effect of CITED1 on ISG expression, the IFNγ vs. dox+IFNγ dataset was reanalyzed using gene set enrichment analysis (GSEA), a statistical tool used to identify phenotypic differences between transcriptome datasets for specific functional gene sets (65). In addition to identifying the top up- and downregulated genes (Fig. 6A), the ‘Interferon gamma response’ gene set was identified as significant among the hallmark gene sets. This produced a negative normalized enrichment score (NES; -1.56), suggestive of an enhanced IFNγ response in CITED1 over-expressing cells (Fig. 6B+C).

**Figure 6:**
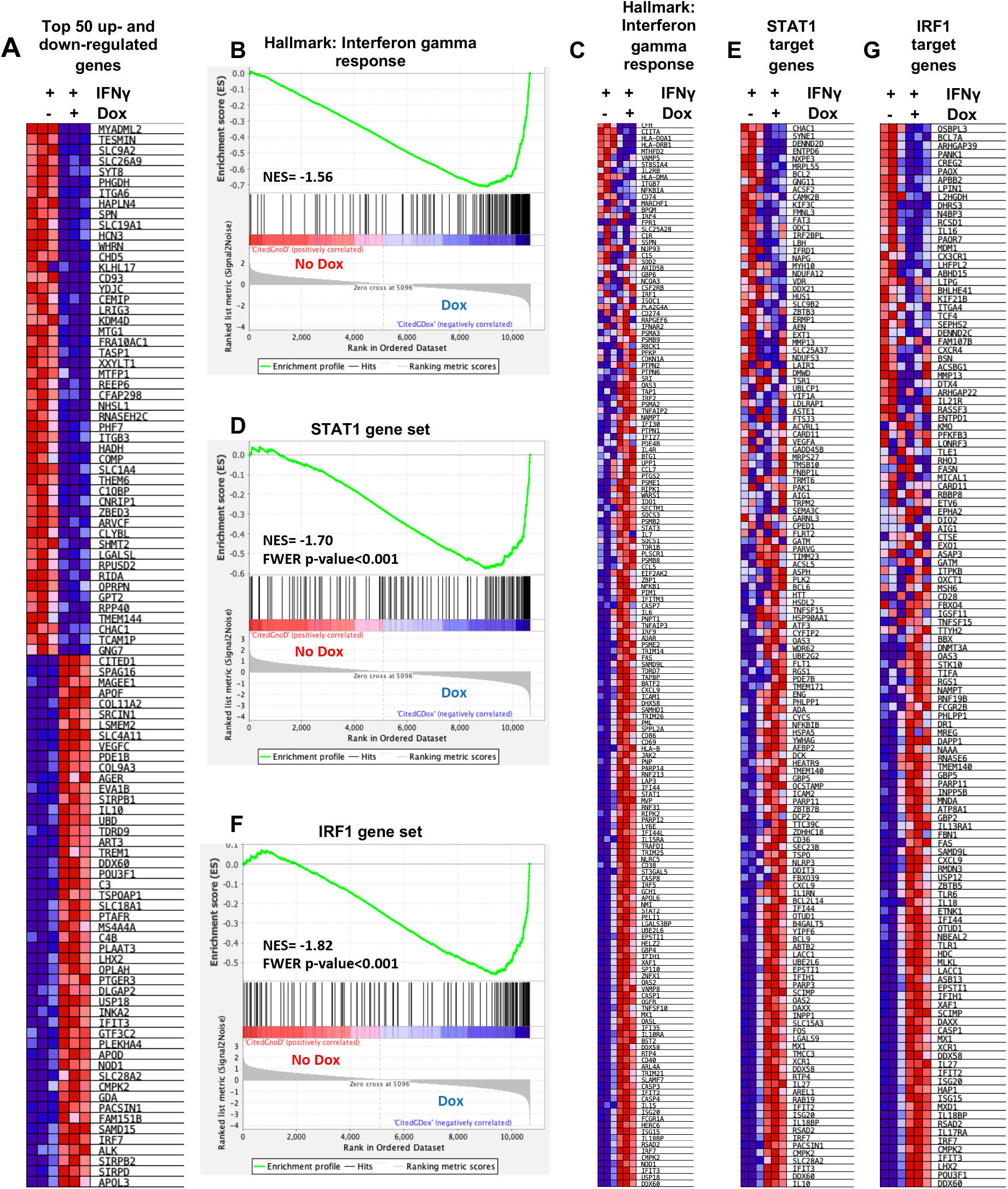
CITED1 modulates the expression of STAT1 and IRF1 target genes. GSEA analysis was performed using the transcriptome profiling dataset described in figure 4A. (A) Heatmap of the top 50 up- and down-regulated genes in IFNγ-stimulated cells with and without dox-induced CITED1 expression. (B+C) GSEA hallmark analysis of the ‘Interferon gamma response’ presented as (B) an enrichment score plot with normalized enrichment score (NES) and (C) heatmap. The same dataset was used together with custom target gene sets to measure the effect on (D+E) STAT1 and (F+G) IRF1 regulated genes and is presented as heatmaps and enrichment score plots. For the analysis using custom target gene sets, a familywise-error rate (FWER) score <0.05 was considered significant. For all heatmaps, red and blue indicate increased and decreased expression, respectively.

As the CuffDiff and GSEA analysis identified genes containing ISRE cis-regulatory sites, such as *Ifit3, Mx1*, and *Isg15* ((21,66,67)), as well as genes containing both or composite ISRE and GAS sites, including *Bst2* (22,24,68), it was plausible that CITED1 affected both STAT1 and IRF1-dependent signaling. This was tested computationally in GSEA using custom STAT1- and IRF1-regulated genes lists. Genes included in these lists (provided by the Mahbeleshwar lab, Case Western University, (44)) were identified based on gene expression changes observed in STAT1 and IRF1 KO macrophages following IFNγ treatment (69,70). This analysis indicated that CITED1 over-expression disproportionately enhanced the expression of genes identified as regulated by STAT1 (Fig. 6D+E) and IRF1 (Fig. 6F+G). This is consistent with the notion that CITED1 and CITED2 have opposing effects on ISG expression as CITED2 null BMDMs show increased expression of STAT1 and IRF1 target genes (44).

### CITED1 knockout attenuated the transcription response to IFNγ

As our gain-of-function experiments indicated that CITED1 functions as a positive regulator of select ISGs, corresponding loss-of-function experiments were performed to verify this result. For these, a RAW264.7 cell line with a null *Cited1* allele was produced using CRISPR/Cas9 and gRNA targeting exon 3 of the gene (Fig. 7A). As RAW264.7 cells are derived from male BALB/c mice, they contain a single copy of the X-chromosome-encoded *Cited1* gene. Therefore, INDELs produced by CRISPR gene-editing were hemizygous. These were characterized by sequencing a 1.5 kb region surround the sgRNA binding site (Fig. 7B). Of the 17 clones screened, 9 contained frameshifts resulting in premature stop codons. Clones 6 and 9 both showed a loss of *Cited1* mRNA and protein expression, as measured by multiplex RT-PCR and western blot, respectively (Fig. 7C+D).

**Figure 7:**
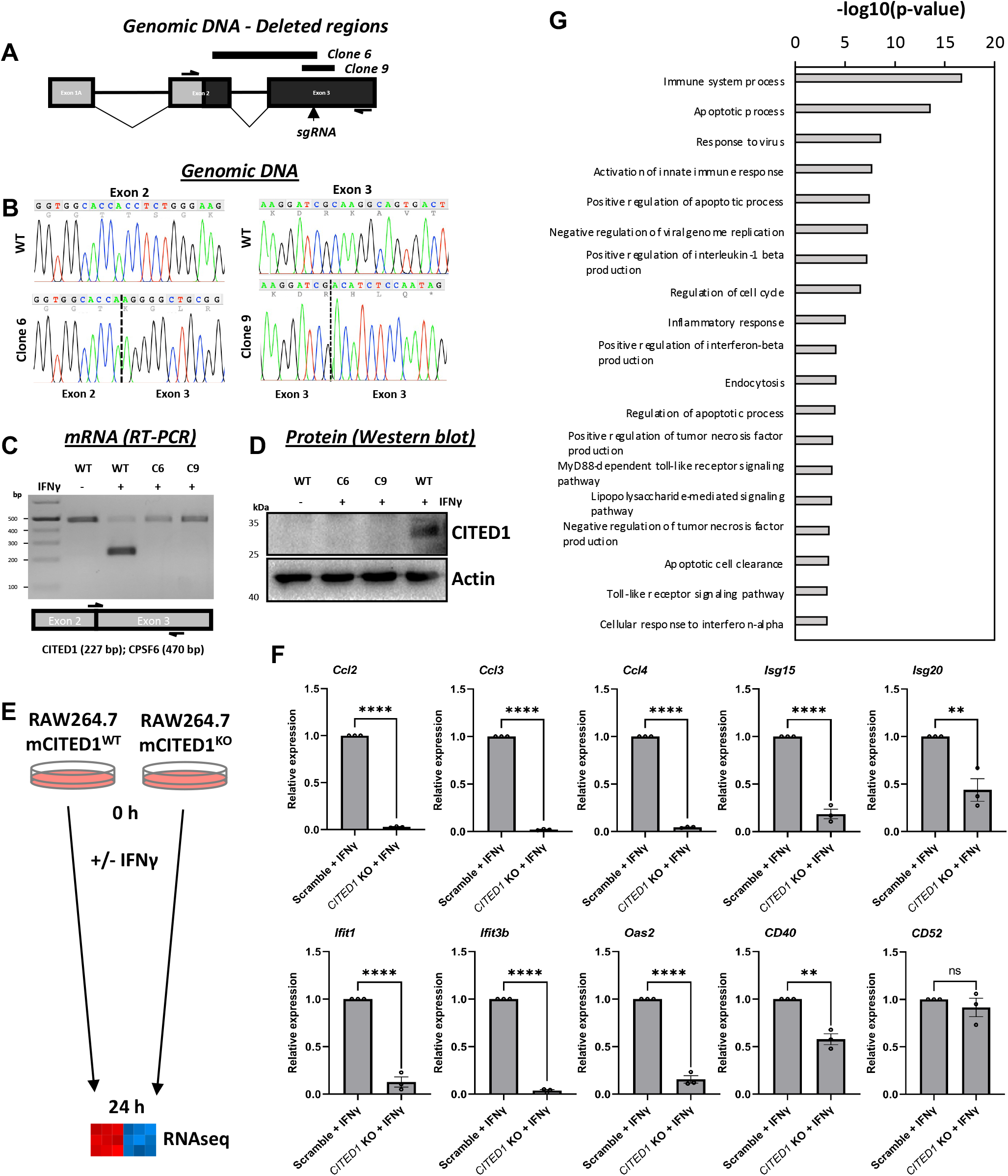
Downregulation of ISGs in CITED1 KO cells. (A) Map of murine *Cited1* gene structure showing protein coding regions of exons (solid black) and the region targeted by an sgRNA to produce KO cells (vertical arrow). The region between the horizontal primer arrows was amplified and subjected to Sanger sequencing to characterize INDELs in edited clonal cells. Deletions resulting in frame shifts were identified in clones 6 and 9 (C6, and C9). The regions of genomic DNA deleted for both clones are represented by the solid black lines above the gene map. (B) DNA chromatographs covering the edited region of the *Cited1* gene in clones 6 and 9 (bottom row) and wild type cells (top row). (C+D) To confirm loss of both *Cited1* gene expression, CITED1 KO clones and wild type RAW264.7 control cells were incubated +/-IFNγ for 24 h. (C) *Cited1* mRNA expression was measured by multiplex RT-PCR using primers for *Cited1* and *CPSF6* as an internal control. The position of *Cited1* forward and reverse primers relative to the exon 2-3 boundary is marked in the diagram below the gel image. (D) CITED1 protein expression was measured by western blotting. (E) For the loss-of-function transcriptome profiling experiments, RAW264.7 scramble control and CITED1 KO cells were incubated +/-IFNγ for 48 h and total RNA was harvested for RNAseq. For the experiment described in (E), a selection of DEGs were validated by qRT-PCR, and (G) GO analysis was performed in DAVID. A selection of the top BP GO terms are displayed ranked by -log(*p*-value).

To appraise the effect of CITED1 loss on ISG expression, the transcriptome of CITED1 null cells (clone 9) was compared to RAW264.7 cells stably expressing Cas9 and a non-targeting scramble gRNA at 48 h post-IFNγ (Fig. 7E). This time was selected based on our timecourse western blot data showing heightened CITED1 expression in unmodified RAW264.7 cells at this time (Fig. 1C). As expected, loss of CITED1 expression reversed the increased expression of numerous ISGs observed in CITED1 over-expressing cells (Table 1), including the 10 genes featured in Fig. 4B. To verify this, result the change in mRNA levels between IFNγ-treated scramble control and CITED1 null cells for this same 10 gene set were measured by qRT-PCR. Here, a statistically significant decrease in expression was seen in the CITED1 null cells for all genes except *Cd52* (Fig. 7F). GO analysis of this data set identified BP terms that were consistent with the macrophage response to IFNγ (Fig. 7G), many of which were common with the corresponding analysis performed for the CITED1 over-expression experiment (Fig. 4C). These included ‘response to virus’, ‘inflammatory response’, and ‘positive regulation of tumor necrosis factor production’. Viewed collectively, these data indicate that over-expression and loss of CITED1 are impacting an overlapping set of genes, suggesting that it has a genuine role in the regulation of the transcriptional response to IFNγ and functions primarily as a positive regulator of genes regulated by STAT1 and IRF1 in this context.

To confirm that loss of CITED1 expression negatively impacted the overall transcriptional response to IFNγ, the same transcriptome dataset was reanalyzed using GSEA. Consistent with our initial analysis, a positive NES score was reported for hallmark ‘Interferon gamma response’ gene set, indicating that this phenotype was more closely associated with the control than the CITED1 null cells (Fig. 8A+B). Positive NES scores were also obtained for analysis using the custom STAT1 (Fig. 8C+D) and IRF1 gene lists (Fig. 8E+F), the opposite to that observed for the corresponding gain-of-function experiments (Fig. 6).

**Figure 8:**
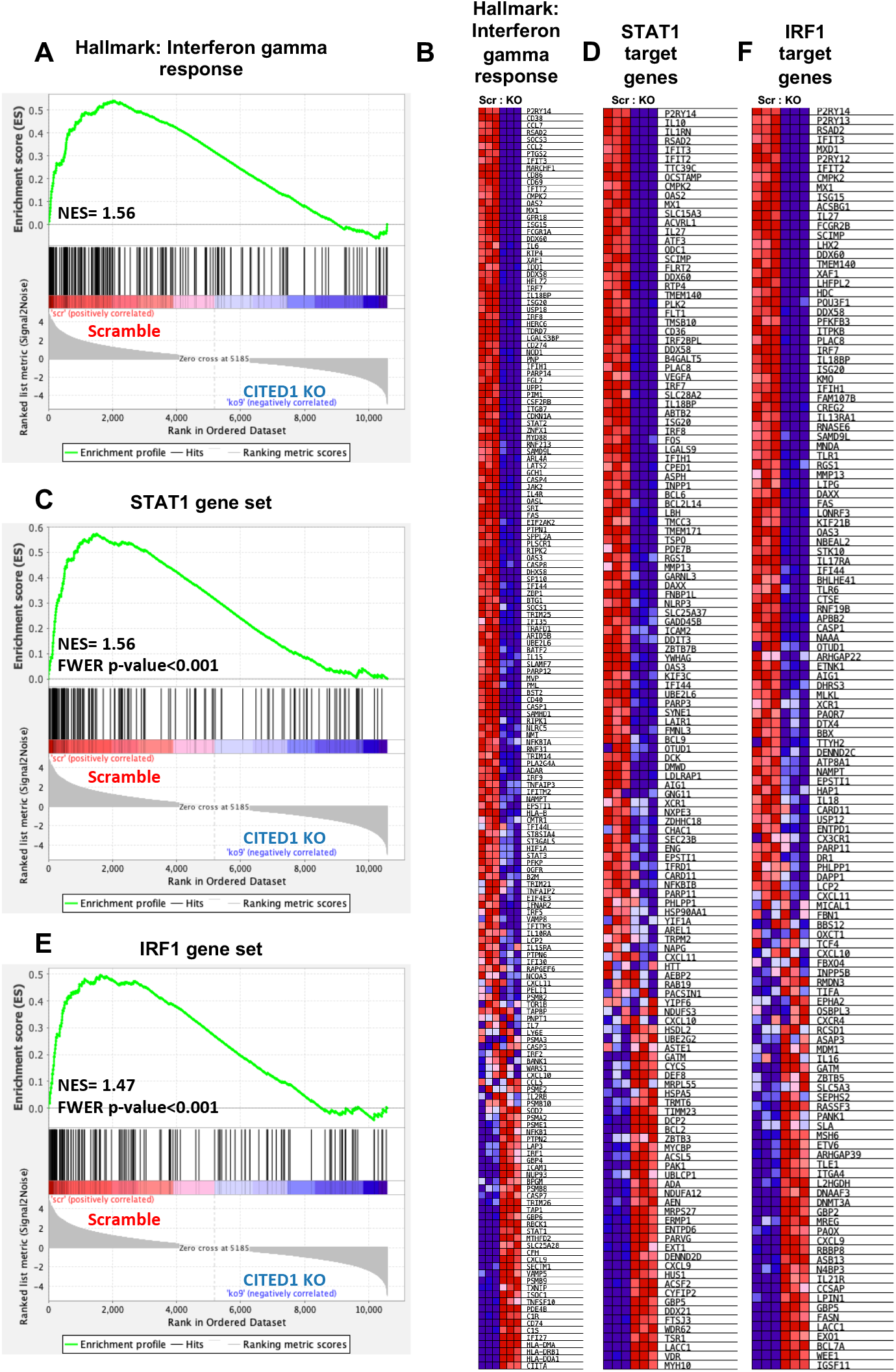
Loss of CITED1 negatively affects STAT1 and IRF1 target gene expression. GSEA analysis was performed using the transcriptome profiling dataset described in figure 7E. (A+B) GSEA hallmark analysis of the ‘Interferon gamma response’ presented as (A) an enrichment score plot with normalized enrichment score (NES) and (B) heatmap. The same dataset was used together with custom target gene sets to measure the effect on (C+D) STAT1 and (E+F) IRF1 regulated genes and is presented as heatmaps and enrichment score plots. For the analysis using custom target gene sets, a familywise-error rate (FWER) score <0.05 was considered significant. For all heatmaps, red and blue indicate increased and decreased expression, respectively.

## Discussion

In this study, we show for the first time that *Cited1* is an IFNγ-responsive gene (Fig. 1A+B) with CITED1 proteins functioning as positive regulators of select ISGs (Fig. 9). In this regard, it is functionally distinct from the closely related CITED2, a well-characterized suppressor of myeloid proinflammatory gene expression (32,39-42,44). This antagonistic relationship between members of transcription factor and co-regulator families is relatively common, with this case being somewhat reminiscent of that between the Kruppel-like Factor (KLF) family of transcriptional factors, which also participate in the control of macrophage-mediated inflammation. Amongst these, Klf2 is largely anti-inflammatory, while Klf6 enhances macrophage proinflammatory gene expression (71,72).

**Figure 9:**
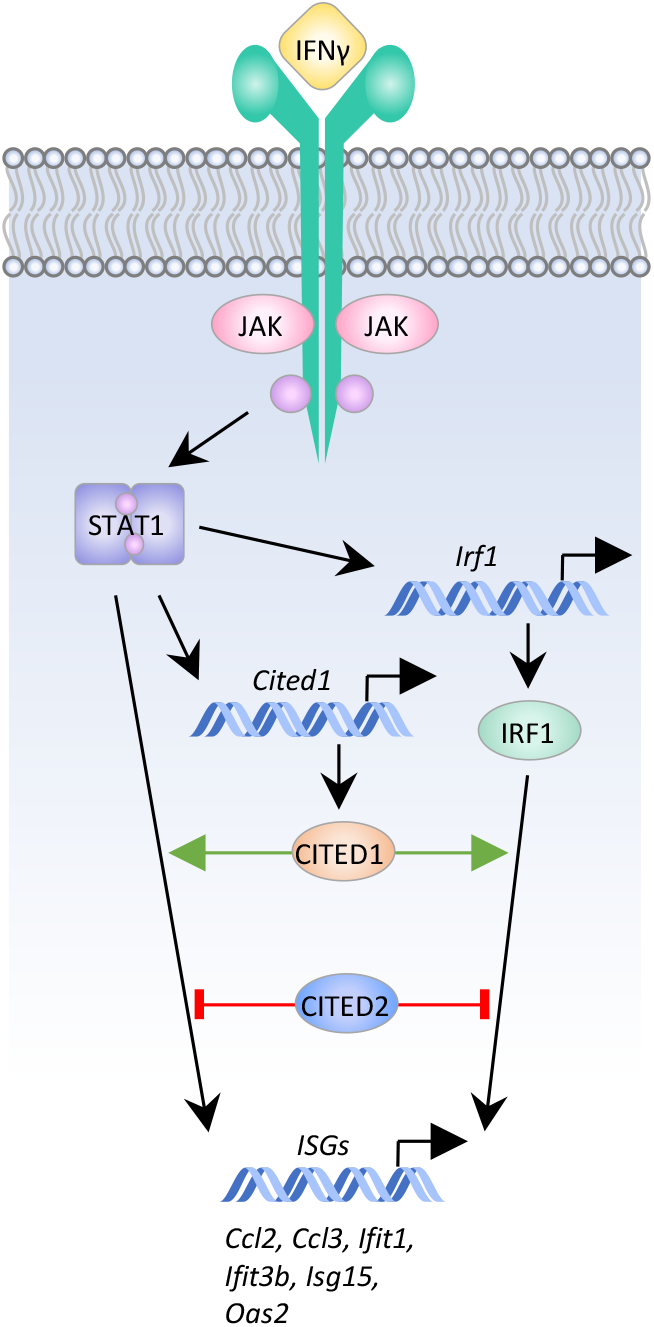
CITED1 as a regulator of ISG expression. IFNγ-stimulated activation of STAT1 promotes expression of IRF1. STAT1 and IRF1 operate in concert to increase the expression of numerous ISGs. Data from the current study shows that IFNγ-stimulation also increases expression of *Cited1* in a STAT1-dependent manner. Unlike CITED2, which represses ISG expression, CITED1 proteins increase the expression of select ISGs.

While the mechanism driving the contrasting effects of these two CITED proteins is currently unclear, it likely stems from their differing interactions with CBP/p300. CITED2 suppresses the activity of proinflammatory transcriptional regulators, including the STATs, p65 NF-κB, and HIF-1α (32,41,42,44), by preventing these proteins from recruiting CBP/p300 to cis-regulatory sites. Here, it acts as a competitive inhibitor, interacting with the N-terminal CH1 domain of CBP/p300, the same region used for docking with these transcription factors (38,39,73,74). While CITED1 also associates with the CH1 domain in *in vitro* immunoprecipitation assays, it shows a strong preference for the CH2 domain, located within a central portion of CBP/p300 that contains HAT activity (37). While this has not been tested, it is conceivable that CITED1 binding via CH2 may still permit the formation of CBP/p300 complexes with proinflammatory transcription factors that dock via CH1. If this is the case, it may even stabilize these complexes by simultaneously interacting with both proteins or stimulating conformational changes in CBP/p300 that enhance transcription factor binding. As an alternative explanation, a feedback relationship between *Cited1* and *2*, as is seen for other gene families (e.g., feedback between p53 and transactivation-deficient isoforms of its homologs, p63 and p73; (75,76)), could at least partially account for these results. For example, inhibition of *Cited2* gene expression or CITED2 CBP/p300 binding by CITED1 could indirectly result in enhanced ISG expression. However, these possibilities were not supported by the data; no change in *Cited2* expression was detected in our transcriptome profiling studies, and increased CITED2 protein expression was not detected in the CITED1 null cells (data not shown).

In this study, we also show that *Cited1* and *2* exhibit contrasting responses to proinflammatory stimuli. *Cited1* is transcriptionally silent in the absence of stimuli and is only expressed in cells incubated with IFNγ for ≥24 h (Fig. 1A+B). In this way, *Cited1* does not participate in the regulation of basal STAT1-regulated gene expression and its delayed expression prevents it from affecting the early phases of an IFNγ response. Rather, it enhances the expression of IFNγ-responsive genes at later timepoints. By contrast, we (and others (32)) show that *Cited2* is constitutively expressed in macrophages (Fig. 1D), and likely functions to suppress inappropriate proinflammatory gene expression in the absence of stimulus and limit it in the presence of persistent inflammatory signals. While there is disagreement in the literature concerning the effects of proinflammatory stimuli on *Cited2* expression (32,41), these data support a model where *Cited2* is transiently down-regulated within 6 h post-IFNγ, possibly to disinhibit or permit the initial phases of STAT1- and IRF1-directed gene expression (Fig. 1D).

Regarding the regulation of *Cited1* itself, our data clearly show that it is downstream of STAT1. Computational promoter analysis indicates that the *Cited1* promoter contains both STAT1 and IRF1 cis-regulatory sites (Fig. 2B), which suggests several plausible models for its regulation; it could be regulated by either transcription factor independently, or it could belong to a large subset of ISGs, including *Gbp2, Lmp2*, and *Socs1*, that are co-regulated by both (19,20,77). A comprehensive analysis of ISG promoters has shown that relatively few are regulated by STAT1 alone, with STAT1 binding co-occurring with IRF1 most of the time, although IRF1 binding frequently occurs alone (77). Given that STAT1 drives *Irf1* expression as part of the first wave of ISG transcription (6), co-regulation of *Cited1* by both transcription factors would constitute a coherent feed-forward loop. This mode of regulation serves as a signaling filter for transient stimuli, permitting expression of the target gene only if the activating signal persists for an extended period of time, allowing for the accumulation of both transcriptional regulators. While feed-forward loops generate delays in target gene expression (78), this is unlikely to explain the timing of *Cited1* transcription, which occurs 24-48 h post-IFNγ, as IRF1 is expressed relatively quickly - as part of the first wave of ISG expression - plateauing at ∼6 h post-IFNγ in RAW264.7 (79). This suggests that regulation of the *Cited1* gene is more complex and may require additional changes, such as post-translational modifications of STAT1 and IRF1 proteins, other transcriptional regulators, or changes to the *Cited1* promoter-chromatin context occurring at later times.

In addition to RNA-level control, this study provides evidence that CITED1 is regulated through altered subcellular localization. In contrast to CITED2, which is reportedly a constitutively nuclear protein (41,80), ectopic EYFP-CITED1 proteins are predominantly cytosolic in unstimulated macrophages, but accumulate in the nucleus post-IFNγ. As a transcriptional co-regulator that operates by controlling the interactions of transcription factors with CBP/p300, the cytosolic sequestration of CITED1 in the absence of stimulus may function as an additional check on its activity, much as it does with other transcription factors, such as NF-κB and NF-AT (81,82).

While the mechanisms regulating CITED1 subcellular localization in macrophages are unclear, the leucine-rich nuclear export signal (NES) located in the CR2 domain of CITED1 is likely responsible for the net cytosolic distribution of the protein in unstimulated cells (60). How IFNγ-stimulation promotes the nuclear accumulation of CITED1 is less obvious, but mechanistic data in other systems may prove instructive. In the osteoblastic cell line, MC3T3-E1, PTH stimulation promotes nuclear translocation of CITED1 (62). This requires PKC-dependent phosphorylation of CITED1 on Ser79, which is necessary but not sufficient for nuclear translocation of CITED1. In the current experiments, endogenous CITED1 proteins appeared as at least two bands on western blots (Fig. 1C+2E), suggesting that the protein was also phosphorylated in macrophages. IFNγ also stimulated increased phosphorylation of a pre-existing pool of ectopic CITED1 in these cells. Given that IFNγ also stimulates PKC activity (83,84), it seems plausible that a similar mechanism to that observed in PTH-stimulated osteoblastic cells may be responsible for CITED1 translocation in macrophages. As a potential caveat, Ser79 is not conserved in the human version of the protein, so the overall importance of this site is questionable.

It is also possible that the phosphorylation detected in these studies may regulate CITED1 activity in other ways. In addition to Ser79, murine CITED1 contains five other putative serine phosphorylation sites (Ser17, 67, 69, 73, and 147; NetPhos-3.1 analysis, phosphorylation site scores >0.90), all of which are conserved between the human and mouse CITED1 sequences (human equivalents: Ser16, 63, 67, 71, and 137). Shi *et al* have shown that all five sites must be ablated to prevent the appearance of partially and hyper-phosphorylated CITED1 species on western blots as HEK293 cells transition through S-phase and mitosis, respectively (60). Phospho-mimetic mutants where all five serine residues were mutated to glutamic acid had reduced p300 binding activity and were less able to enhance Smad4-dependent gene expression. This raises the possibility that the phosphorylated CITED1 species detected in the current experiments may also have reduced activity. Future studies mapping the sites of IFNγ-stimulated phosphorylation and measuring the effect of these post-translational modifications on the subcellular localization and ability of the protein to modulate the expression of ISGs will provide clarity on this issue.

In closing, our use of a dual gain- and loss-of-function approach has allowed us to identify candidate CITED1-regulated genes with a high degree of confidence. These include members of gene families that are fundamental to the macrophage IFNγ-response; such as the C-C motif chemokine family genes, *Ccl2* and *Ccl3*, well-characterized regulators of immune cell migration and inflammation (85), but also members of the *Isg* (*Isg15* and *Isg20*) and *Ifit* (*Ifit1* and *Ifit3b*) gene families. Both encode proteins that enhance the anti-viral activity of macrophages, but do so by very different mechanisms. The Ifit proteins act as ubiquitin-like modifiers that restrict viral replication in host cells (86), while ISG proteins enhance macrophage polarization and nitric oxide production in response to viral infection (87). While these data provide important clues to the biological role of CITED1 in innate immune function, a more accurate and holistic view must await future *in vivo* studies. In the meantime, this study points to a greater level of complexity in the control of JAK-STAT signaling and the roles played by the CITED family of proteins in this.

## Materials & Methods

### Mammalian Cell Culture

RAW264.7 macrophage-like cells were obtained from ATCC (Manassas, VA) and were cultured in DMEM supplemented with 10% FBS, 25 mM HEPES, 1% penicillin and streptomycin, and 50 μg/mL gentamicin (all from Sigma Aldrich). Lenti-X 293T packaging cells were obtained from Takara-Clontech and were cultured in DMEM supplemented with 10% tetracycline-free FBS (Takara-Clontech), 200 mM L-glutamine, 1 mM sodium pyruvate, and 100 units/mL penicillin and streptomycin. All cell lines were maintained at 37 ºC in a humidified 5% CO_2_ atmosphere. For all western blotting, qRT-PCR, and RNA sequencing experiments, cells were seeded in 6-well plates (Thermo Fisher Scientific) at a density of 1.5×10^5^ cells/well in 2 mL of medium and grown to ∼80% confluency prior to treatment with the indicated reagents. Unless specified otherwise, recombinant murine IFNγ (Biolegend, San Diego, CA) was used at a final concentration of 200 U/mL, doxycycline at 100 ng/mL, and LPS at 100 ng/mL. For treatments lasting longer than 24 hours, the medium, cytokines, and compounds were replaced at 24-hour intervals.

### Plasmids and lentiviral constructs

The pINDUCER20-mCITED1 lentiviral construct was produced by amplifying the full-length murine CITED1 coding sequence from pCDNA3-Flag-mCITED1 (a kind gift from Dr. Mark P. de Caestecker (VUMC)) and recombining it into the pENTR Gateway donor vector (Addgene, USA) using an In-Fusion ligase-independent cloning kit (Takara-Clontech). The mCITED1 sequence was then subcloned into the pINDUCER20 lentiviral vector (a kind gift from the Weissmiller lab, MTSU; originally sourced from the Elledge Lab; (88)) using an LR Clonase® II reaction (Life Technologies). The pINDUCER20-EYFP-mCITED1 lentiviral construct was produced by amplifying the full-length murine CITED1 coding sequence from pCDNA3-Flag-mCITED1 and subcloning it into pEYFP-C1 (Takara/Clontech) in-frame with the EYFP coding sequence. The complete EYFP-mCITED1 sequence was recombined into the pENTR Gateway donor vector using an In-Fusion ligase-independent cloning kit then subcloned into the pINDUCER20 lentiviral vector. The fidelity of all plasmid constructs was verified using by Sanger sequencing (Eurofins Genomics) prior to use in experiments. Lentiviral constructs encoding the Cas9 enzyme and a gRNA were obtained from VectorBuilder, USA. The sequences (5’ to 3’) of gRNAs targeting murine genes were as follows: *Cited1* TACCCCGGGGTCACCGCAAA, *Stat1* GTTGGGCGGTCCCCCGATGC, and non-targeting “scramble” gRNA GTGTAGTTCGACCATTCGTG.

### Lentiviral transductions

Lentiviral constructs were packaged by transfection in to Lenti-X 293T cells using a Lenti-X™ Packaging Single Shots kit. Viral supernatants were concentrated and tested for the presence of viral particles using Lenti-X GoStix. The concentrated lentiviral supernatants were used to transduce RAW264.7 cells by spinoculation in the presence of polybrene. Transduced cells were selected by growth in 500 μg/mL G418, and then clonal lines were produced by dilution plating. Loss of gene expression was confirmed using a combination of western blotting and RT-PCR. All Lenti-X products were obtained from Takara-Clontech.

### Immunoblotting

For immunoblotting experiments, cells were lysed in RIPA buffer containing protease and phosphatase inhibitors (Sigma). For λ protein phosphatase (λPP) assays, cells were lysed in 1% Triton-X100 buffer instead without phosphatase inhibitors. The protein concentration of the samples was determined by BCA assay and normalized by dilution with the appropriate lysis buffer. To dephosphorylate proteins samples, ∼50 μg of protein were incubated with 20 units of λPP (NEB, USA) for 30 minutes. All samples were boiled in 1× Laemmli buffer and resolved by SDS-PAGE and visualized by western blotting.

### Antibodies

The primary antibodies used for western blotting experiments were as follows: β-actin (A2066, Sigma), MSG-1 (CITED1; D-7, sc-393585, Santa Cruz Biotechnology), and STAT1 (D1K9Y, 14994T, Cell Signaling). Primary antibody binding was detected using mouse anti-rabbit IgG-horseradish peroxidase (HRP; sc-2357, Santa Cruz Biotechnology) or anti-mouse m-IgGkappa binding protein (BP)-HRP (sc-516102, Santa Cruz Biotechnology), as appropriate. Membranes were incubated with standard enhanced chemiluminescent (ECL) reagents or SuperSignal West Atto (Pierce/ThermoFisher), and bands were visualized using a ChemiDoc MP Imaging System with Image Lab Software (Bio-Rad, Hercules, CA).

### RNA extraction and RNA sequencing

Total RNA was extracted using RNeasy Mini Kits (Qiagen), according to the manufacturer’s instructions with genomic DNA removed from samples using a Message Clean kit (GenHunter, Nashville, TN). Clean RNA was resuspended in 10 μL diethyl pyrocarbonate (DEPC)-treated water and shipped to Novogene (Sacramento, CA). Here, RNA quality was appraised using an Agilent 2100 Bioanalyzer and samples with an acceptable RNA integrity number were used for cDNA library production and RNA sequencing using the HiSeq 2500 system to produce 150 bp transcriptome paired-end reads.

### Analysis of RNA sequencing data

Quality checks of fastq RNAseq data files were performed using FastQC (version 0.11.5; (89)). Based on the data quality, no file trimming was necessary. Reads were aligned to the version 38 mouse genome using STAR aligner (version 2.5.3a;(90)). Scaffolding was provided by the mouse reference genome annotation (version 39.90, (91)) in CyVerse Discovery Environment (92). In the Galaxy platform (93), a read count table was generated from the bam output from STAR and the same mouse genome annotation using FeatureCounts (94) and multi-join (95). The resultant read counts were imported into R, and samples were clustered based on their whole genome gene expression profile using EdgeR (93), as described in (96). To evaluate samples for inclusion or exclusion, the data were displayed as a multi-dimensional scale plot. As all samples per condition clustered with their replicate pool (n=3), they were all used to construct transcript annotations. To make pairwise comparisons for differentially expressed genes (DEGs) among replicate groups, CuffDiff2 (version 2.2.1; (97)) was used with the same mouse genome annotation and genome files. From the pairwise comparisons, DEGs with fold change ≥ 2.0 and q ≤ 0.05 were considered biologically relevant and statistically significant. Gene ontology (GO) analysis was performed in Database for Annotation, Visualization, and Integrated Discovery (DAVID) 2021 bioinformatics resource tool (version Dec. 2021; (98,99)). GO terms were ranked and plotted as –log(p-value). For *Cited1* read visualization, bam output files (from STAR), the mouse genome file, and the mouse genome annotation were loaded into the IGV, which was searched for *Cited1* to display its gene structure with read depth.

Gene set enrichment analysis (GSEA) was performed using GSEA software (version 4.2.3; (65,100)). Read counts derived from STAR output (90) were imported into R and the SARTools R package, DESeq2 version (version 1.7.4; (101)) was used to produce normalized read counts per experiment per sample. Per GSEA recommendations for RNA sequencing experiments (65), genes with no counts in any sample were removed, along with low-expressing genes (mean or geometric mean <10 reads across all samples). As recommended in the GSEA user guide, a false discovery rate (FDR) cut of 25% was used for hallmark gene set analysis. STAT1 and IRF1 target gene enrichment analysis was performed in GSEA using custom gene lists assembled by the Mahabeleshwar lab from data published in Langlais *et al* 2016 and Semper *et al* 2014 (69,70), and has previously been used to measure the effects of *Cited2* loss on STAT1- and IRF1-regulated gene expression (44). For this analysis, a family-wise error rate (FWER) <0.05 cut-off was used to determine whether a gene set was significantly enriched.

### RT-PCR

For both reverse transcriptase chain reaction (RT-PCR) and quantitative RT-PCR (qRT-PCR) experiments, RNA (isolated as described above) was reverse transcribed to produce cDNA libraries using Maxima H Minus Reverse Transcriptase in the presence of dNTPs, Oligo dT primers, and RiboLock RNase inhibitor (ThermoScientific). RT-PCR was performed as a multiplex amplification reaction, using primers for *Cited1* and *Cpsf6* as an internal control. PCR products were resolved on 2% agarose gels containing ethidium bromide and visualized under UV illumination. For qRT-PCR the indicated cDNAs of interest were amplified using PerfeCTa SYBR Green FastMix (Quantabio) in a CFX Opus 96 Real-Time PCR Instrument (Bio-Rad).

Normalization was performed using two reference genes (*Cyc1* and *Actb*). Primers were designed such that at least one primer for each pair spanned an exon-exon boundary and produced a product size between 70 and 150 bp. The sequences (5’ to 3’) of primer pairs used were as follows: *Actb* F-CACTGTCGAGTCGCGTCC, R-TCATCCATGGCGAACTGGTG, *Ccl2:* F-CAGATGCAGTTAACGCCCCA, R-TGAGCTTGGTGACAAAAACTACAG, *Ccl3:* F-CCAAGTCTTCTCAGCGCCATA, R-TCTCTTAGTCAGGAAAATGACACC, *Ccl4:* F-CTGTGCAAACCTAACCCCGA, R-AGGGTCAGAGCCCATTGGT, *Cd40:* F-TTGTTGACAGCGGTCCATCT, R-TTCCTGGCTGGCACAAATCA, *Cd52:* F-CAAAGCTGCTACAGAGCCCA, R-CCAAGGATCCTGTTTGTATCTGAAT, *Cited1:* F-CTGCCACCGATTTATCGGACTT, R-CTCCTGGTTGGCATCCTCCTT, *Cited2:* F-GCAAAGACGGAAGGACTGGA, R-CGTAGTGTATGTGCTCGCCC, *Cpsf6:* F-TTACACTGGGAAGAGAATCGC, R-CTGGAAAAGGTGGAGGTGG, *Cyc1* F-CTAACCCTGAGGCTGCAAGA, R-GCCAGTGAGCAGGGAAAATA. *Ifit1:* F-TCTGCTCTGCTGAAAACCCA, R-CACCATCAGCATTCTCTCCCAT, *Ifit3b:* F-CCTTCCTGCCAAGGATTGCT, R-TGTGATCAAAAGGTGGTCTGTGA, *Isg15:* F-TCTGACTGTGAGAGCAAGCAG, R-CCTTTAGGTCCCAGGCCATT, *Isg20:* F-TGAAGCCAGGCTAGAGATCC, R-AGGGCATTGAAGTCGTGCTT, and *Oas2:* F-GCCTTGGAAAGTGCCAGTACC, R-CCTTGGTCCTGCCACAAGAT.

### Genotype-Tissue Expression Analysis

The online Geneotype-Tissue Expression (GTEx) resource was employed to compare the expression of human *Cited* family members across 54 non-diseased tissues from ∼1000 individuals. The following Ensembl gene IDs were used for the analysis ENSG00000125931 (*Cited1*), ENSG00000164442 (*Cited2*), and ENSG00000179862 (*Cited4*).

### Promoter analysis

Putative transcription factor binding sites in the *Mus musculus Cited1* promoter were identified using the Eukaryotic Promoter Database (EPD; Swiss Institute of Bioinformatics) with the Jasper core 2018 vertebrates transcription factor motif library (56). The promoter (NCBI Reference Sequence: NM_007709) was scanned from -2000 to 100 bp relative to the transcriptional start site with a cut-off (p-value) of 0.001.

### Fluorescence Microscopy

Clonal RAW264.7 cells stably transduced with pINDUCER20-EYFP-CITED1 were plated at 1.5×10^5^/dish with 2 mL medium in 35 mm glass-bottom dishes (Cellvis, USA) and incubated with 100 ng/mL doxycycline to induce EYFP-CITED1 protein expression prior to IFNγ treatment for the indicated periods. Cells were stained with 2.5 μg/mL Hoechst 33342 30 min prior to imaging using a Zeiss LSM700 confocal laser scanning microscope equipped with a Plan-Apochromat 63x magnification/1.40 numerical aperture oil immersion DIC M27 objective lens and controlled using Zen software (Zeiss). EYFP fluorescence was excited using a 488 nm laser. Hoechst 33342 was excited using a 405 nm laser and detected through a 490 nm short-pass filter. To calculate nuclear:cytoplasmic EYFP-CITED1 ratios, mean nuclear and cytoplasmic fluorescence intensity measurements were made using Fiji (102), and these data were exported to Excel.

### Statistical analysis

All experiments were performed as three discrete biological repeats unless otherwise stated. Statistical analyses were performed in GraphPad Prism 7 (GraphPad, USA) using the tests indicated in the figure legends.

## Availability of data and materials

All data generated or analyzed during this study are included in this published article. All non-commercial plasmid constructs used in this study are available on request from David E. Nelson.

## Acknowledgements

We thank the Mahabeleshwar lab (Case Western Reserve University) for providing assistance with the GSEA analysis using STAT1 and IRF1 target gene sets. Thanks also to the de Caestecker lab (Vanderbilt University) for providing the mCITED1 expression plasmid.

## Author contributions

DEN conceived, designed, and coordinated this study, wrote the manuscript, and performed the experiments shown in Fig. 2E, 7B+C, and the analysis in Fig. 2B, 4C-D, 5, and 7G. RLST assisted with the design of the RNAseq assays and performed the bioinformatic analysis of the data used for Fig. 1A+G, 4-8, and Tables 1+2. E.E.M. assisted with the design and coordination of the study. AS performed and analyzed the experiments shown in Fig. 1A-F, 2C, 4B, 7D+F, and prepared all samples used for RNAseq-based transcriptome profiling. MH performed and analyzed the experiments shown in Fig. 3. HK performed and analyzed the experiment shown in Fig. 2C. GEM assisted with the preparation of samples for the experiment described in Fig. 4. All authors reviewed the results and approved the final version of the manuscript.

## Consent for publication

Not applicable.

## Ethics approval and consent to participate

Not applicable.

